# *Pseudomonas* taxonomic and functional microdiversity in the wheat rhizosphere is cultivar-dependent and links to disease resistance profile and root diameter

**DOI:** 10.1101/2024.06.13.598955

**Authors:** Courtney Horn Herms, Rosanna Catherine Hennessy, Frederik Bak, Ying Guan, Patrick Denis Browne, Tue Kjærgaard Nielsen, Lars Hestbjerg Hansen, Dorte Bodin Dresbøll, Mette Haubjerg Nicolaisen

## Abstract

Diversity within lower taxonomic units in microbial communities is a key trait, giving rise to important ecological functions. In the rhizosphere, these functions include disease suppression and pathogen inhibition. However, limited effort has been given to exploring intragenus microdiversity in an increasingly homogenous agricultural system. Through an integrative approach combining culture-dependent and -independent methods, we explore the rhizosphere *Pseudomonas* pangenome and demonstrate cultivar-dependent taxonomic and functional microdiversity between two closely related modern winter wheat cultivars. A *Fusarium-*resistant cultivar demonstrated increased *Pseudomonas* taxonomic diversity but not biosynthetic diversity when compared to the susceptible cultivar, coinciding with a thinner root diameter of the resistant cultivar. We found enrichment of *Pseudomonas* isolates capable of antagonizing *Fusarium* as well as chitinase-encoding genes and pyoverdine gene clusters in the resistant cultivar. Across closely related *Pseudomonas* isolates from the two cultures, there were differences in genomic content and biosynthetic gene clusters. Ultimately, we highlight the need for fine-scale analysis to uncover the hidden microdiversity within rhizosphere *Pseudomonas*.

## 1. Introduction

Plants and soil microorganisms interact through such a tight collaboration that the plant and its rhizosphere microorganisms are considered a holobiont (Vandenkoornhuyse et al., 2015). The microbiome in the rhizosphere generally assembles into a stable community comprised primarily of the phyla Actinobacteria, Bacteroidetes, Firmicutes, and Proteobacteria, which harbor key functions important for plant health (Bai et al., 2022). But, at the order and genus level, plant microbiome communities can differ because of plant type, root exudation, and age (Qu et al., 2020), leading to functional community differences between closely related crops (Mendes et al., 2018).

Diversity within genera and species, i.e., microdiversity (Moore et al., 1998), plays an important role in microbial ecology (García-García et al., 2019). Bacterial species arise when populations split into ecologically distinct fractions allowing for specific niche adaptations (Koeppel et al., 2013). In this case, genes involved in functions such as iron acquisition and biofilm formation become part of lineage-specific gene pools (Shalev et al., 2021). Thus, increased diversity within genera and species contributes to ecological interactions through resource and niche competition as well as direct antagonism (Jousset et al., 2011; Hu et al., 2016; Yang et al., 2017; Li et al., 2023; Spragge et al., 2023).

Despite this, only few studies to date have addressed microdiversity of rhizosphere microbiomes (Mauchline et al., 2015; Oni et al., 2019; Pacheco-Moreno et al., 2021, 2024). These studies demonstrate the potential genomic and function diversity yet to be discovered in the rhizosphere environment. The lack of fine taxonomic resolution in microbiome studies of the plant holobiont may be a consequence of short-read amplicon-based analyses that conceal diversity and functional differences between closely related microbes (Chiniquy et al., 2021; Edgar, 2018; Wang et al., 2022). Advances in sequencing technologies are increasing amplicon read lengths, allowing for better culture-independent identification of soil microbes (Stevens et al., 2023), but isolation efforts, whole genome sequencing, and pangenome analysis remain the gold standard for the identification of microdiversity (Huang et al., 2023; Zhang et al., 2024).

The small body of research on rhizosphere microdiversity has focused on comparing cultivars distinct by their breeding history or cropping strategies, but there is limited attention to closely related modern cultivars that demonstrate their own microdiversity in terms of root exudation and root morphology (Iannucci et al., 2021). Root exudation is a plant trait that is key to contributing to microbial community diversity (Feng et al., 2023; Sasse et al., 2018; Yang et al., 2023), but it alone cannot fully explain rhizosphere microbiome assembly (Takamatsu et al., 2023). The physical landscape is also suggested as a driver of microbial community diversity (Dubey et al., 2021). The physical and consequent chemical landscapes implicated by different root phenotypes have the potential to modulate root-microbe interactions (Herms et al., 2022). Specifically, there is growing evidence for a link between thinner root diameter and bacterial diversity in the rhizosphere for both woody and herbaceous species (Fleishman et al., 2023; King et al., 2023; Luo et al., 2021; Pervaiz et al., 2020; Saleem et al., 2018; Zai et al., 2021). However, to date, this relationship has not been examined in cereals, nor have microbial microdiversity and root diameter been related.

A genus of particular interest for the study of microdiversity in the rhizosphere is *Pseudomonas*. This ubiquitous soil genus boasts high intrinsic intragenus genomic diversity (Girard et al., 2021; Loper et al., 2012; Lopes et al., 2018) and a wide assortment of secondary metabolites (Gavriilidou et al., 2022; Gross and Loper, 2009; Gu et al., 2020). Additionally, *Pseudomonas* is a key member of disease-resistant microbiomes (Hong et al., 2023; Lv et al., 2023; Qiu et al., 2022) thanks to their production of a wide spectrum of bioactive secondary metabolites such as siderophores, phenazines, cyclic lipopeptides (CLPs), and chitinases (Mauchline and Malone, 2017; Pacheco-Moreno et al., 2021).

In the present work, we explored the root-associated *Pseudomonas* taxonomic and functional microdiversity of closely related and commercially available modern cultivars of winter wheat (*Triticum aestivum* L.) grown in agricultural soil. By choosing two cultivars with contrasting *Fusarium culmorum* resistance profiles: the resistant Sheriff cultivar and the susceptible Heerup cultivar (Jørgensen et al., 2023, 2020), we further aimed at investigating whether a link between microdiversity and cultivar resistance to soil-borne fungal infections could be observed. We undertook an integrative approach combining culture-dependent and culture-independent pangenome analyses coupled to root morphological analysis. We expected the taxonomic and functional microdiversity to be different between the two cultivars. Specifically, we hypothesized that the *Fusarium*-resistant cultivar, Sheriff, harbors a more taxonomically and functionally diverse soil *Pseudomonas* community. Second, we hypothesized that pseudomonads from the Sheriff cultivar inhibit *F. culmorum* growth and harbor a greater abundance of genes that may be responsible for antagonizing *F. culmorum*. Lastly, we hypothesized that the expected increase in Sheriff *Pseudomonas* microdiversity would coincide with a smaller root diameter of the Sheriff root system.

## 2. Materials and Methods

### 2.1 Wheat cultivation and sampling

Two winter wheat (*Triticum aestivum* L.) cultivars, Heerup and Sheriff, were chosen based on their documented resistance to *F. culmorum* in the field and were supplied by Sejet Plant Breeding (Horsens, Denmark). One seed of each cultivar was sown in a 24 x 7 cm PVC pot filled with soil mixed with sand (DANSAND, Denmark; filter sand no. 2) in a 2:1 ratio. The soil was a sandy loam collected from the plough layer (0-25 cm) in the University of Copenhagen’s Long Term Nutrient Depletion Experiment located at Højbakkegård in Høje Taastrup, Denmark (van der Bom et al., 2017). The soil had received mineral NPK fertilizer at a rate of 120 kg nitrogen, 20 kg phosphorus and 120 kg potassium ha^-1^y^-1^ for the past 25 years and contained 170 g kg^−1^ clay, 174 g kg^−1^ silt, 336 g kg^−1^ fine sand, 255 g kg^−1^ coarse sand, and 40 g kg^−1^ organic matter. The soil had a pH(CaCl2) of 5.4, an Olsen-P content of 11.4 mg kg^−1^. The water holding capacity (WHC) was 33%. The soil was air-dried and sieved to 8mm prior to the experiment.

The plants were grown under controlled greenhouse conditions in three stages: an 18-day germination period (19°C day/15°C night; 16h day/8h night; light intensity 300 µE), followed by a 13- week vernalization period (6°C day/4°C night; 8h day/16h night; light intensity 150 µE), and finally a 17-week growth period to maturity (19°C day/15°C night; 16h day/8h night; light intensity 500 µE). Pots were regularly rotated and watered to 70% WHC by weighing.

Three different time points were sampled from three independent setups following the above method (Methods Fig 1). After four weeks, a set of plants were sampled for root morphology scanning. A second set of plants were sampled for *Pseudomonas* isolation at first node emergence (T1). A third set of plants were used for two rounds of long-read 16S rRNA amplicon sequencing, one at first node emergence (T1) and one at flag leaf emergence (T2).

**Figure 1.**
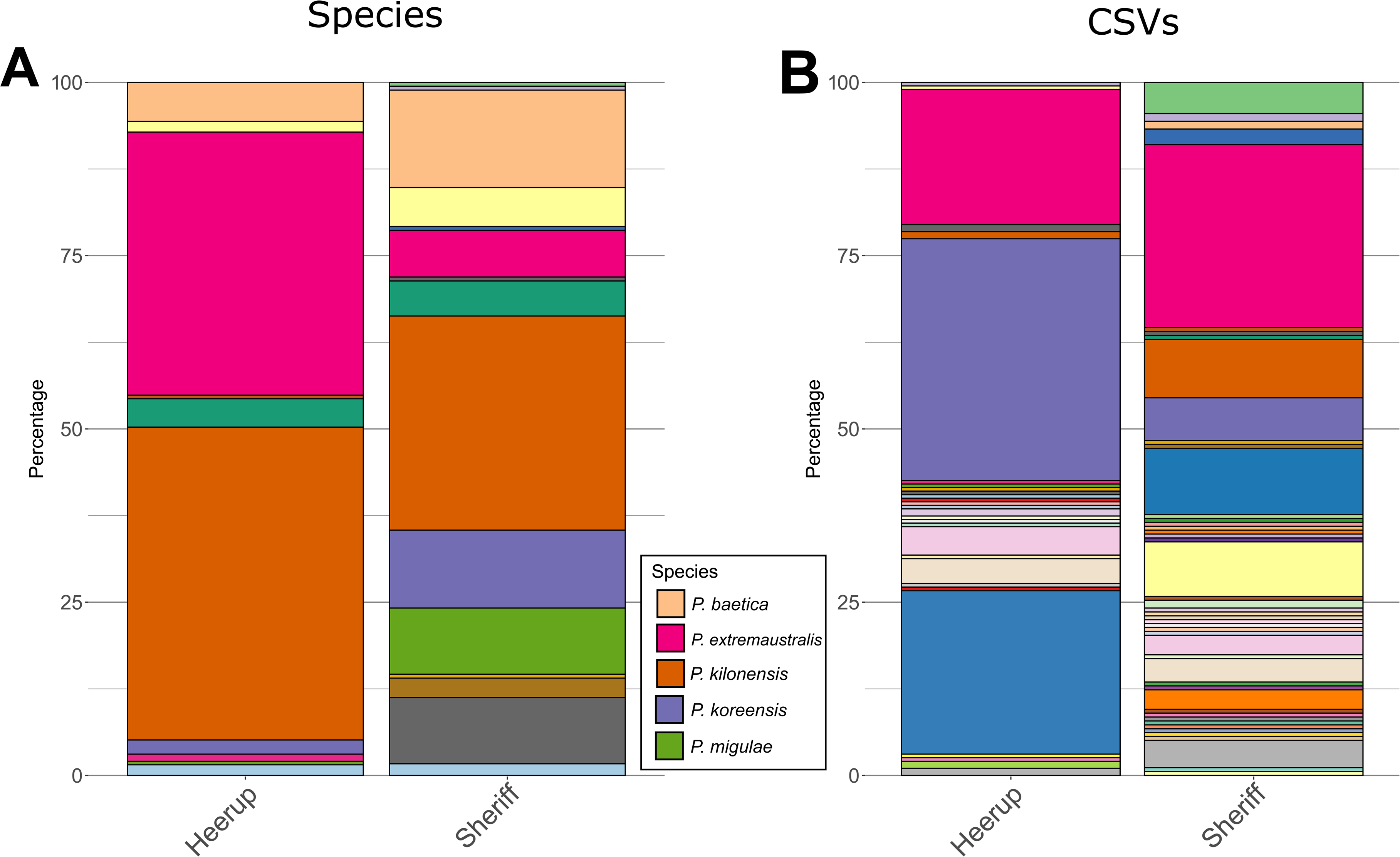
Composition of the Heerup and Sheriff Pseudomonas strain libraries identified by Sanger sequencing of minimum 900 bp of the 16S rRNA gene from 195 purified isolates from Heerup and 178 purified isolates from Sheriff. Each color represents a Pseudomonas species or CSV. A) Composition at the species level identified by sequence alignment to the NBCI 16S rRNA gene database. A legend identifying the top five most abundant species is shown. B) Composition at the CSV level, defined as 100% identity across a minimum of 900 bp of the 16S rRNA gene.

### 2.2 *Pseudomonas* library building

Three replicate plants were sampled at first node emergence (T1) to obtain *Pseudomonas* colonies (Methods Fig 1). To obtain rhizoplane samples, one gram (corresponding to approximately 4 cm) of root was washed in 25 mL sterile PBST (10% 10x PBS, 0.1% Tween 80) and subsequently placed in 25 mL of fresh, sterile PBST. The roots were sonicated in a water bath for two minutes to release rhizoplane soil and microorganisms. To isolate *Pseudomonas* species, 50 μL of undiluted and a 1:10 dilution in PBST (PBS, 0.1% Tween 80) of the rhizoplane sample was plated onto Gould’s S1 media (Gould et al., 1985) to select for *Pseudomonas* isolates. After incubation at room temperature for two to four days, single colonies were streaked for purification first on Gould’s S1 media and a second time on Luria-Bertani (LB) agar (Miller, 1972). Pure isolates were maintained as frozen stocks (15% glycerol w/w).

### 2.3 16S rRNA gene sequencing of cultured isolates

A modified version of colony PCR (von Stein et al., 1997) was used to amplify the full-length 16S rRNA gene of the isolates in the strain library. A 30 μL PCR reaction mixture was prepared: 15 μL Master Mix (Supreme NZYTaq II 2x Master Mix, NZYTech, Portugal), 0.6 μL of 10 μM forward and reverse primer (27F, 1492R; <u>Weisburg et al., 1991)</u>, 11.8 μL Sigma water, and 2 μL 1:1000 diluted overnight culture. The cycling conditions were as follows: preliminary boiling step of 96°C for 5 min followed by 25 cycles of denaturing at 98°C for 10 s, annealing at 57°C for 30 s, extension at 72°C for 45 s, and a final extension of 10 min at 72°C. Successful amplification of the 16S rRNA gene was visualized on 1% agarose gels. PCR purification and Sanger sequencing of the PCR product was completed by Eurofins (Germany). The ends of the reads were trimmed by Eurofins based on quality scores. Reads of short length (<900 base pairs (bp)) were manually discarded using CLC Genomics Workbench (v. 12.0.03) (QIAGEN).

Sequences were oriented to the same strand using USEARCH (Edgar, 2010) and the RDP training set (v. 18) as reference and then aligned using MUSCLE (Edgar, 2004). Gaps at the end of the aligned sequences were removed using Gblocks (Castresana, 2000). Sequences were clustered into sequence variant cluster of 100% identity using the USEARCH *cluster_fast* command (Edgar, 2010). The 16S rRNA sequences from each sequence variant cluster were submitted to NBCI BLASTn for alignment to the 16S rRNA database (Altschul et al., 1997).

The 16S rRNA genes of the strain library isolates are available in Genbank under accession numbers PP894319-PP894691.

### 2.4 Phenotypic assays

The wheat fungal pathogen *F. culmorum* was sporulated in Mung bean media (Ilmi et al., 2019) (20 grams boiled mung beans, strained, in 1 L distilled water). A high-throughput, agar-based confrontation assay was developed to screen the strain library for antifungal activity. Briefly, the *Pseudomonas* isolates were inoculated onto a pre-germinated lawn of 1.45 x 10^3^ *F. culmorum* spores on 1/5 potato dextrose agar (BD DIFCO, Denmark). After three days of co-inoculation at room temperature, the isolates were scored for their ability or inability to inhibit *F. culmorum* growth. A positive result required a clearance zone of at least 1 mm around a colony. *Serratia inhibens* S40 (Hennessy et al., 2020) was used as the positive control.

A high throughput drop collapse assay (Bodour and Miller-Maier, 1998) was used to screen the strain library for biosurfactant production. The isolates were grown in 1.5 mL liquid LB media for four days in a deep-well 96-well plate at 28°C while shaking (200 rpm). To test for biosurfactant production, 40 μL of culture was pipetted onto Parafilm, and at least two minutes passed before the droplet size was measured. Water was used as the negative control, and *P. fluorescens* SBW25, which produces the surfactant viscosin (Alsohim et al., 2014), was used as the positive control.

### 2.5 Genome sequencing and bioinformatics pipeline

A representative isolate from each unique culture sequence variant (CSV) clusters identified from the strain library via 16S rRNA gene sequencing were subject to full genome sequencing. CSV clusters containing many isolates were sequenced deeper, proportional to the size of the CSV cluster and how many isolates in the cluster originated from each cultivar. Genomic DNA was prepared from overnight cultures using the QIAGEN Genomic DNA Handbook using the QIAGEN Buffer Kit (Germany) or the Genomic Mini Ax Bacteria 96-Well kit (A&A Biotechnology, Poland) according to the manufacturer’s instructions. Quality was checked using a NanoDrop ND-1000 spectrophotometer (Thermo Fisher Scientific, Carlsbad, CA, USA) prior to library building. Genomic DNA was prepared for sequencing using the Rapid Barcoding Sequencing Kit (SQK-RBK004) or the Rapid Sequencing gDNA barcoding kit (SQK-RBK110.96) and sequenced on MinION and PromethION flow cells (v. 9.4.1) (Oxford Nanopore Technologies, United Kingdom). Reads were basecalled with Guppy (v. 6.2.1) using the “super accuracy” dna_r9.4.1_450bps_sup.cfg basecalling model. Adapter sequence and barcodes were trimmed from basecalled reads with Porechop (v. 0.2.4). Genomes were assembled using Flye (v. 2.9-b1774) (Lin et al., 2016; Kolmogorov et al., 2019), polished with Medaka (v. 1.7.1) (https://nanoporetech.github.io/medaka), and annotated using Prokka (v. 1.14.6) (Seemann, 2014) using default parameters. The quality of the genomes was assessed using CheckM (v1.1.10) (Parks et al., 2015). Genomes not passing thresholds for completeness (> 90.0%) and contamination (< 2.0%) were removed from future analyses. Dereplication with dRep (v. 3.4.5) (Olm et al., 2017) and the Genome Taxonomy Database Toolkit (v. 2.3.0) (Chaumeil et al., 2020; Parks et al., 2020) were used to delineate and identify species and strains. In dRep, 95% average nucleotide identity (ANI) was chosen to delineate species, and 99.55% ANI was used to delineate strains.

The annotated genomes were screened for secondary metabolite prediction with antiSMASH (v. 7.0) (Medema et al., 2011) using the standard cutoff of 0.3, *glocal* mode, and the *include_singletons* flag. BiG-SCAPE (v. 1.1.7) (Navarro-Muñoz et al., 2020) and the MiBIG database (v. 3.1) (Terlouw et al., 2023) were used for biosynthetic gene cluster family analysis. Anvi’o (v. 7.0) (Eren et al., 2021) was used to assign functions using NCBI’s Clusters of Orthologous Groups (COGs) database (Tatusov et al., 2000), generate a pangenome, and quantify functional enrichment between sets of genomes (Shaiber et al., 2020).

The fully assembled genomes are available under BioProject accession number PRJNA1108081.

### 2.6 Long-read 16S rRNA gene amplicon sequencing

For long-read 16S rRNA amplicon sequencing, rhizoplane samples were obtained as described above at first node (T1) (n = 5 for Heerup and Sheriff) and flag leaf emergence (T2) (n = 10 for Heerup and n = 9 for Sheriff) (Methods Fig 1). After the sonication step, the rhizoplane samples were flash frozen in liquid nitrogen and freeze-dried. DNA was extracted using the FastPrep-24 5G bead-beating system (MP Biomedicals, Irvine, CA, USA) at 6.0 m/s for 40 s and the FastDNA SPIN Kit for soil (MP Biomedicals) according to the manufacturer’s instructions.

PCR reactions were prepared with 32.5 μL Sigma water, 10 μL SuperFi Buffer (Thermo Fischer), 0.5 μL Platinum SuperFi DNA Polymerase (Thermo Fischer), 1 μL 10 mM dNTPs, 2.5 μL each of 10 nM forward and reverse primer (TAG Copenhagen, Denmark), and 1 μL DNA template diluted to 5 ng/μL. *Pseudomonas*-specific primers for amplifying 969 bp of the 16S rRNA gene (position 289-1258) were used (forward primer 5’-GGTCTGAGAGGATGATCAGT-3’ and reverse primer 5’- TTAGCTCCACCTCGCGGC-3’) (Widmer et al., 1998). ZymoBIOMICS Microbial Community DNA was used as a sequencing control (Zymo Research, Irvine, CA, USA). The cycling conditions were as follows: preliminary denaturation step of 98°C for 30 s followed by 30 cycles of denaturing at 98°C for 10 s, annealing at 63°C for 30 s, extension at 72°C for 45 s, and a final extension of 10 min at 72°C. Successful amplification was visualized on 1% agarose gels. PCR products were purified using AMPure XP beads at a 0.6 ratio (Beckman Coulter Inc., Brea, CA, USA). Amplicons were prepared for sequencing using a native barcoding kit (Oxford Nanopore SQK-NBK114.96) and sequenced on a Minion (Oxford Nanopore) on an R10.4.1 flow cell (Oxford Nanopore FLO-MIN114). Raw nanopore data was basecalled with Guppy (v. 6.3.9) using the “super accuracy” basecalling model dna_r10.4.1_e8.2_260bps_sup-v4.1.0.

Adapters were trimmed from Nanopore reads using porechop (v. 0.2.4) (https://github.com/rrwick/Porechop) with the following options: adapter_threshold 95, extra_end_trim 0, end_threshold 95, middle_threshold 95, extra_middle_trim_good_side 0, extra_middle_trim_bad_side 0, min_split_read_size 800. The options were used to filter out overly short inserts and to leave the primer sequences in the trimmed reads. Following this, primer sequences were identified, and the intervening sequences (inserts) were extracted using strip_degen_primer_deep (github.com/padbr/asat) and inserts between 880 and 970 nt were retained. High quality inserts (Phred quality ≥ 30) were identified using Nanofilt (De Coster et al., 2018). High quality inserts were then dereplicated and unique sequences counted using the fastx_uniques utility of usearch (v. 11.0.667) (Edgar, 2010), and zero-width amplicon sequencing variants (ASVs) were inferred using unoise3 (Edgar, 2016). All read inserts trimmed of adapters and primers, within the 880 to 970 nt range, but not quality filtered, were used to create a ASV table using the otutab utility of usearch.

The raw reads are available under BioProject accession number PRJNA1108081.

### 2.7 Root morphology scanning

To measure root morphology of the two cultivars, 4-week old plants were removed from the pots and all soil was rinsed from the roots using tap water (n = 9 for Heerup and n = 10 for Sheriff).

Samples of 1 cm length were taken 1 cm (top), 11 cm (middle), and 18 cm (bottom) from the top of each root (Methods Fig. 1). Images of each section of root were photo-scanned using the Epson Perfection V700 scanner (8-bit gray scale, 600 dpi), and root diameter was measured using RhizoVision (v. 2.0.3) using “broken roots” analysis mode (Seethepalli et al., 2021). To remove debris and artifacts from the analysis, a maximum object size was set at 10 mm ^2^, and edge smoothing was set at a threshold of three. After segmentation, root pruning was performed with a threshold of 50. Root diameter ranged were manually set at 0-0.2 mm, 0.2-0.3 mm, 0.3-0.4 mm, 0.4-0.5 mm, 0.5-1.0 mm, and 1.0-1.5 mm. After RhizoVision analysis, the percentage of the root system in each of diameter class of the three root sections were averaged to obtain a single percentage for each plant replicate.

### 2.8 Statistical analysis

All statistics were performed in R (v. 4.3.1). The rarefaction plot was generated using vegan (v. 2.6-4) (Oksanen et al., 2022). Calculating the effect of cultivar on antifungal activity, biosurfactant production, and unique isolates was performed by Fischer’s Exact test. When comparing means, all data were tested for normality using the Shapiro-Wilk test. The two-sided Wilcoxon Rank Sum test was used in place of a two-sided unpaired Student T test when normality could not be assumed. P values were adjusted for multiple testing as appropriate using the Benjamini & Hochberg method (Benjamini and Hochberg, 1995). Plots were made using ggplot2 (v. 3.4.2).

Long-read amplicon sequencing data were analyzed using Qiime2 (Caporaso et al., 2010). Normalization of the ASV table was done by rarefying with all ASVs. Following this, the ASV table was filtered to keep only features classified in the *Pseudomonas* genus. Qiime2 was then used to derive values for α-diversity (richness, Chao1, and Shannon) and β-diversity (Bray-Curtis dissimilarity). The Shannon diversity values were converted to effective Shannon diversity by raising two to the power of the Shannon diversity. Composition data and α-diversity metrics were compared between groups of samples using the Mann-Whitney U-test to test for significance. ASVs were omitted from composition tests if the ASV is not seen in at least 50% of the samples in one of the groups. Anosim (Clarke, 1993), with 999 permutations, was used to test if two or more groups of samples differed from each other.

## 3. Results

### 3.1 Culture-based screening indicates a more diverse and antifungal Sheriff *Pseudomonas* microbiome

A strain library of 373 pseudomonads comprising 195 isolates from Heerup and 178 isolates from Sheriff was successfully isolated. Sanger sequencing of the 16S rRNA gene confirmed all 373 isolates as *Pseudomonas*. Clustering at 100% identity across a minimum of 900 bp and an average of 1114 bp of the 16S rRNA gene resulted in 65 unique culture sequence variant (CSV) clusters, and BLASTn alignment to the NBCI 16S rRNA gene database revealed 17 potential *Pseudomonas* species (Table S1). The most abundant species in the library was identified as *P. kilonensis* with 143 isolates, followed by *P. extremaustralis* and *P. baetica* with 86 and 36 isolates, respectively. Three and seven species and 17 and 36 CSVs were uniquely recruited to the Heerup and Sheriff cultivar, respectively (Figure 1). Additionally, the Sheriff cultivar harbored significantly more CSVs than the Heerup cultivar; 33 CSVs were found in Heerup and 59 CSVs in Sheriff (Fisher’s exact test, *p* = 0.0052).

However, the rarefaction curve demonstrated that the full *Pseudomonas* diversity was not sampled via isolation (Figure S1).

To assess functional microdiversity within the isolated *Pseudomonas*, a high-throughput antifungal screen and drop-collapse assay were performed. Across both cultivars, 33% (n = 129) of strains inhibited growth of the fungal pathogen *F. culmorum*, whereas 17% (n = 67) of strains were positive for biosurfactant activity. The high-throughput phenotypic screens showed that antifungal isolates were enriched in the Sheriff strain library (40%) compared to Heerup strain library (27%) (Fisher’s exact test, *p* = 0.0030), while the proportion of biosurfactant producers did not differ between Sheriff (20%) and Heerup (14%) (Fisher’s exact test, *p* = 0.0603) (Fig. 2).

**Figure 2.**
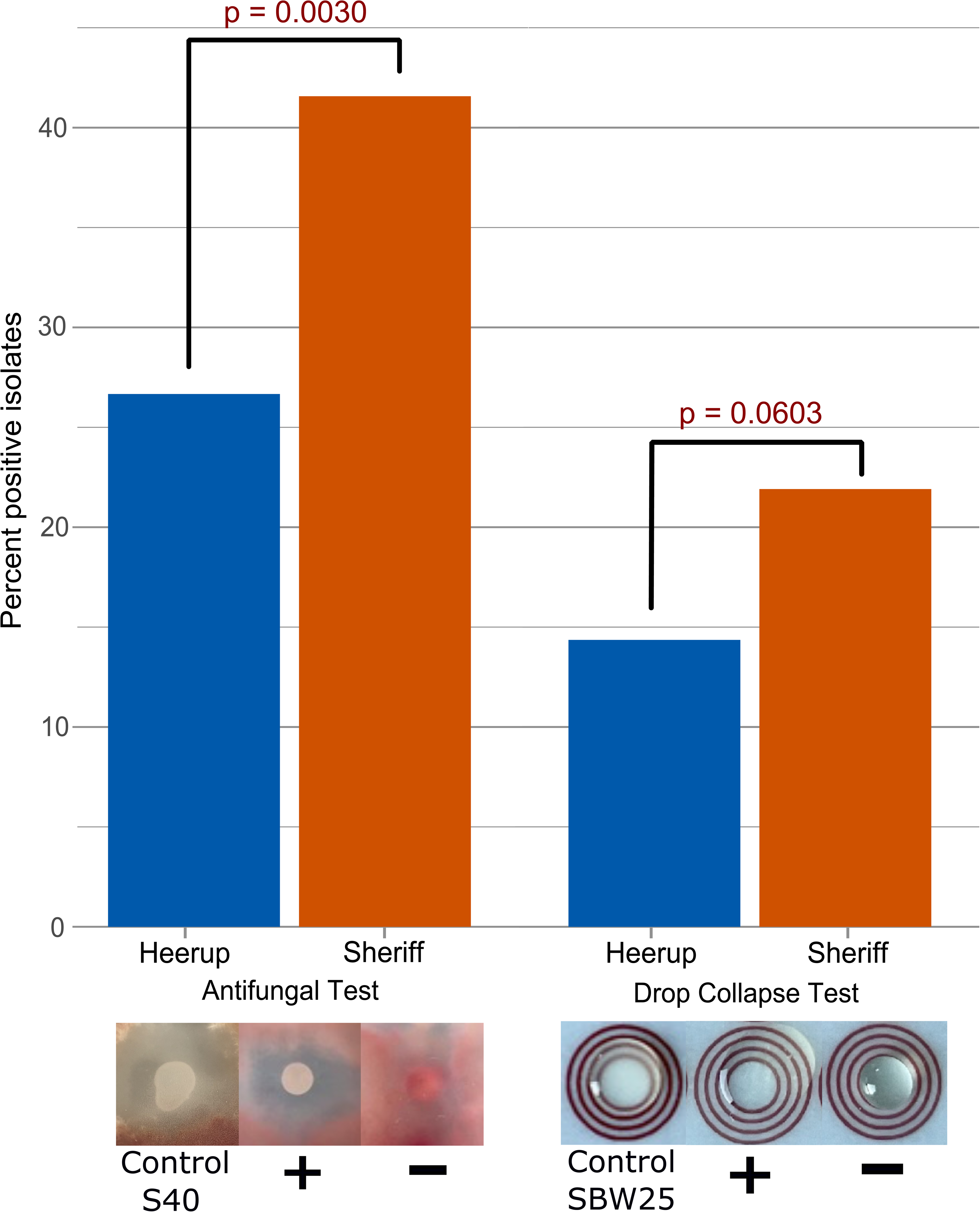
In vitro F. culmorum inhibition and biosurfactant production of the Heerup and Sheriff strain libraries. P-values of Fisher’s exact test are given above the bars. Strains Serratia inhibens S40 and Pseudomonas fluorescens SBW25 were used as the positive control for the antifungal assay and the biosurfactant assay, respectively.

### 3.2 Targeted long-read 16S rRNA amplicon sequencing supports *Pseudomonas* microdiversity and antifungal potential in Sheriff cultivar

To confirm Sheriff’s enrichment of *Pseudomonas* microdiversity in a culture-independent manner, long-read 16S rRNA amplicon sequencing of the rhizoplane sample was performed on plants from a separate experiment from the culturing. Sampling was performed at the same time point as the culturing (at first node emergence, T1) (Heerup and Sheriff, n = 5) and one month after the culturing time point (at flag leaf emergence, T2) (Heerup n = 10, Sheriff n = 9).

Prior to rarefying, 373,093 reads were assigned to 707 ASVs, with a range of 2,014 to 37,612 reads per sample. Seventy of the 707 ASVs were classified within the *Pseudomonas* genus. Exact matches to the 16S rRNA genes of the 65 unique CSVs were found in the amplicon data, but due to the length of the amplicon at 880-970 bp, many 16S rRNA gene sequences from the CSVs were grouped together into one ASV. Twenty-two additional, uncultured *Pseudomonas* ASVs were identified.

However, the full *Pseudomonas* diversity and richness was still not captured with amplicon sequencing, especially at T2 (Fig S2A-B).

At T1, 67% to 91% of reads per sample were classified as *Pseudomonas* with a median of 81% (Fig S2C). One uncultured pseudomonad, ASV7, dominated all samples at this time point and contributed to a low percentage of cultured *Pseudomonas* reads captured by sequencing (Fig S2D). There was also no significant difference in *Pseudomonas* diversity nor any ASVs differentially abundant between the two cultivars at T1 (data not shown).

At T2, ASV7 decreased 95-fold in relative abundance, coinciding with a drop in *Pseudomonas* reads per sample, with a range of 2% to 43% with a median of 5% (Fig S2C) and an increased percentage of cultured *Pseudomonas* reads (Fig S2D).

At T2, Sheriff demonstrated a higher diversity of *Pseudomonas* than Heerup (Mann-Whitney U-test, *p* = 0.00013), which aligned with the conclusion from culturing. One uncultured ASV and three ASVs that were exact sequence matches to the 16S rRNA sequence of five isolates were significantly enriched in the Sheriff cultivar (Table 1A). Within the ASVs matching to cultured isolates that were enriched in the Sheriff cultivar at T2, two of the isolates showed antifungal activity against *F. culmorum in vitro*.

Across T1 and T2, three ASVs that were exact sequence matches to the 16S rRNA sequence of 12 isolates were significantly enriched in the Sheriff cultivar compared to the Heerup cultivar (Table 1B). Within the ASVs matching to cultured isolates that were enriched in the Sheriff cultivar across T1 and T2, seven of the isolates were antifungal against *F. culmorum in vitro.* At no time point were there any ASVs enriched in the Heerup cultivar.

**Table 1A.** Enriched ASVs in Heerup and Sheriff at T2, with notice of the isolates that were antifungal in the *in vitro* screen. P values were determined using the Mann-Whitney U-test. ASVs were omitted from composition tests if the ASV is not seen in at least 50% of the samples in one of the groups.

| ASV | Antifungal isolates<br>within ASV | median number of<br>reads in Sheriff | median number of<br>reads in Heerup | Pval |
| --- | --- | --- | --- | --- |
| S1E05 | 0/1 | 1.0 | 0.0 | 0.0472 |
| S2B12_S2C03 | 1/2 | 3.0 | 0.0 | 0.0322 |
| ASV67 | n/a | 1.0 | 0.0 | 0.0098 |
| H3E07_S3A09 | 1/2 | 1.0 | 0.0 | 0.0398 |

**Table 1B.** Enriched ASVs in Heerup and Sheriff across T1 and T2, with notice of the isolates that were antifungal in the *in vitro* screen. P values were determined using the Mann-Whitney U-test. ASVs were omitted from composition tests if the ASV is not seen in at least 50% of the samples in one of the groups.

| ASV | Antifungal<br>isolates within<br>ASV | median number<br>of reads in<br>Sheriff | median<br>number<br>of reads<br>in Heerup | Pval |
| --- | --- | --- | --- | --- |
| H1D09_H2B05_H2D02_S1A03_S2<br>D10_S3C01_S3E11_S3F07_S3H10 | 6/9 | 14.0 | 6.0 | 0.0305 |
| H3E07_S3A09 | 1/2 | 1.0 | 0.0 | 0.0036 |
| H3H08 | 0/1 | 1.0 | 0.0 | 0.0079 |

### 3.3 Genome sequencing highlights *Pseudomonas* microdiversity and identifies chitinase as potential driver for enriched *Pseudomonas* antifungal activity in Sheriff cultivar

For a more detailed analysis on functional microdiversity, isolates were chosen for full-genome sequencing based on 16S rRNA gene phylogeny. Briefly, a representative isolate of each of the 65 unique CSVs was sequenced, and large CSV clusters were sequenced deeper, proportional to the size of the CSV cluster and how many isolates in the CSV cluster originated from each cultivar. A total of 112 isolates were fully genome sequenced of which 53 were isolated from Heerup and 59 were isolated from Sheriff.

Dereplication (Olm et al., 2017) and the Genome Taxonomy Database (Chaumeil et al., 2020) identified 21 species and 48 strains in the library (Table S1). Six species and 14 strains were unique to Heerup while 10 species and 31 strains were unique to Sheriff. Hence, only five species were isolates from both cultivars. Gene clusters generated based on amino acid sequence similarity were generated using Anvi’o (Eren et al., 2021). In total, 14,769 gene clusters present in at least two genomes were identified. The core genome, defined as genes present in all 112 genomes, consisted of 2,322 gene clusters.

Gene cluster analysis revealed differences in genomic content among isolates within the same *Pseudomonas* species (Fig 3A). This genomic mirodiversity was, in some instances, influenced by cultivar of origin. In the case of the six isolates identified as *P. brassicacearum* strain R, three loci composed of 65 gene clusters, 6 gene clusters, and 9 gene clusters, respectively, were present only in the four isolates from the Heerup cultivar while absent in the two isolates from the Sheriff cultivar (Fig 3B). In this same taxon, a locus of 24 gene clusters was present only in the Sheriff-isolated strains (Fig 3B); this locus was predicted to be associated with the synthesis of flagella (Table S2).

**Figure 3.**
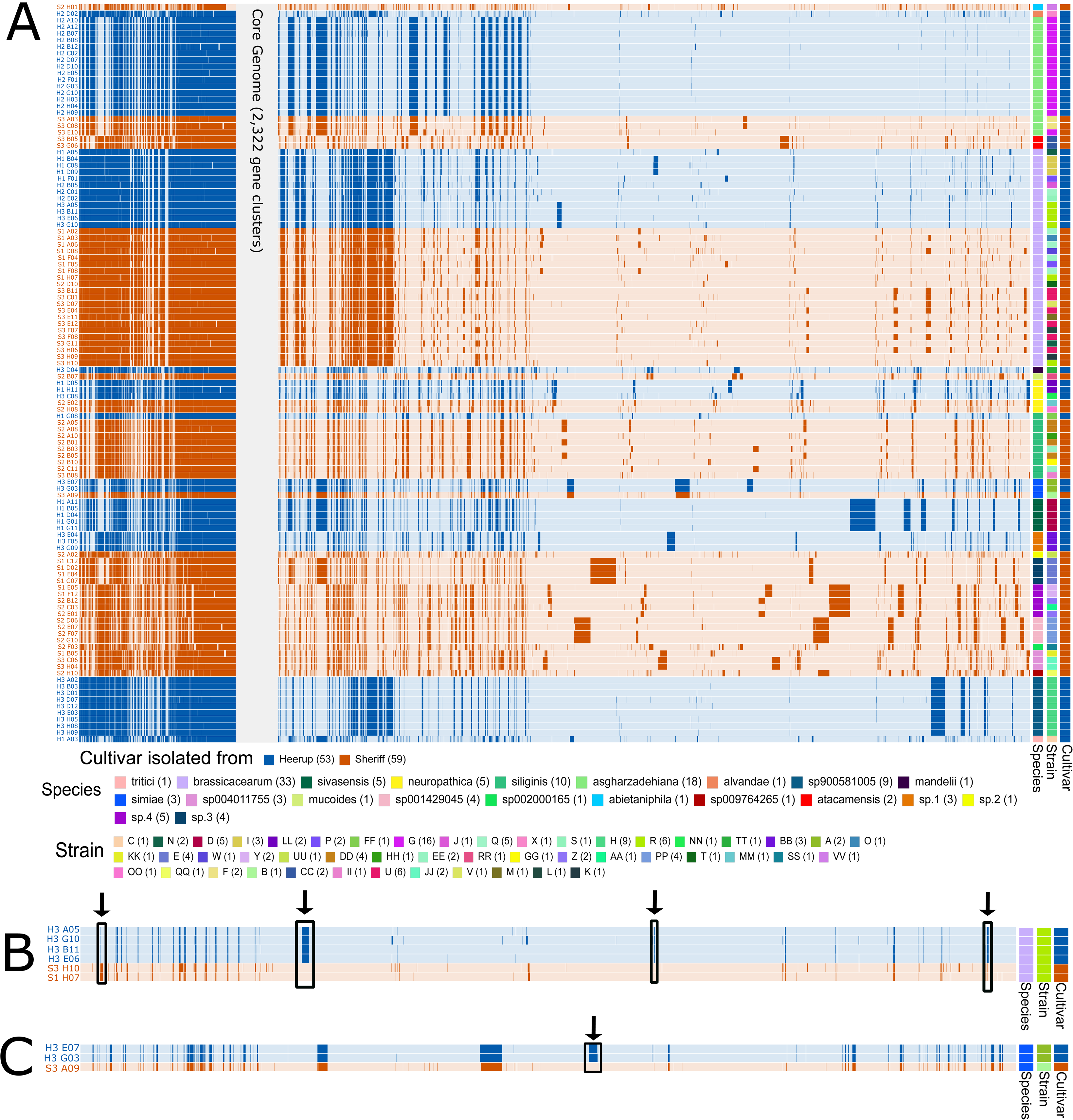
Pangenome of the 112 genomes. A) Blue highlight represents an isolate cultured from Heerup, and an orange highlight represents an isolate cultured from Sheriff. Shaded bars indicate the presence of gene clusters in the genome. The core genome of 2,322 gene clusters is collapsed for increased clarity. Species and strain identification is indicated. B) Highlight boxes and arrows indicates the missing gene cluster locus in the Heerup-isolated strains of P. brassicacearum strain R, and the missing three gene cluster loci in Sheriff-isolated strains, despite its presence in closely related (same species and strain) isolates. C) Highlight box and arrow indicates the missing one gene cluster locus in Sheriff-isolated strains of P. simiae despite its presence in closely related (same species) isolates from Heerup.

Additionally, there is a locus of 76 gene clusters in *P. simiae* that were only present in isolates cultured from the Heerup cultivar (Fig 3C).

In order to link genomic microdiversity to functional potential, enrichment scores for COG20 pathways, categories, and functions were calculated between isolates cultured from Heerup and Sheriff using Anvi’o (Eren et al., 2021). There were no differentially enriched COG20 pathways. Twenty-nine COG20 categories were differentially enriched between strains originating from the two cultivars (adjusted *q*-value < 0.05): 15 in Heerup and 14 in Sheriff (Table S3). The most common differentially abundant COG20 categories between the two cultivars were signal transduction mechanisms (n = 10), secondary metabolites biosynthesis, transport and catabolism (n = 7), and cell membrane biogenesis (n = 7). A total of 213 COG20 functions were differentially enriched between strains originating from the two cultivars (adjusted *q*-value < 0.05): 104 in Heerup and 109 in Sheriff (Table S4). Notably, chitinase family GH19 (accession: COG3179) was enriched in isolates originating from the Sheriff cultivar (adjusted *q*-value = 9.2E-6). Of the remaining differentially enriched COG20 functions, no others were known to be directly implicated in antifungal activity (Table S3).

### 3.4 Genome mining reveals *Pseudomonas* functional partitioning between Heerup and Sheriff via cultivar-specific species enrichment

We mined the genomes for biosynthetic gene clusters (BGCs) to examine the biosynthetic potential of the *Pseudomonas* strain library. Screening with antiSMASH (Medema et al., 2011) and analysis via BiG-SCAPE (Navarro-Muñoz et al., 2020) revealed 1315 BGCs in the strain library that grouped into 137 BGC families of 21 product classes. The most common class of BGC in the strain library was ribosomally synthesized and post-translationally modified peptides (RiPP, n = 450) followed by NRPS (n = 378). The most common product class was RiPP-like (n = 287), followed by redox co-factors (n = 118), NRPS-like (n = 117), and NRPS (n = 111). The BGC families identified in the strain library were mapped to the MiBIG database, and this revealed six known products (Table 2).

**Table 2.** BGC families in the strain library that were identified as known products and the cultivar they were enriched in, if applicable. Significant differences in the abundance of the BGC family between cultivars was calculated using the two-sided Wilcoxon Rank Sum test with fdr-adjustment using the Benjamini & Hochberg method. Significant p-values (p < 0.05) are bolded.

| BGC family | Known BGC | Product Class | Enriched in | Adjusted p-value |
| --- | --- | --- | --- | --- |
| 2428 | 3-thiaglutamate | RiPP-like | n/a | 0.22 |
| 2549 | Viscosin | NRPS | n/a | 0.09 |
| 2653 | Arthrofactin A, anikasin, lokisin, stechlisin B2 | NRPS | n/a | 0.84 |
| 3049 | Pf-5 pyoverdine | NRPS | Sheriff | **0.0297** |
| 3453 | 2,4-diacetylphloroglucinol | T3PKS | n/a | 0.37 |
| 3764 | Tabtoxin | blactam | n/a | 0.36 |

To test the hypothesis of increased functional diversity and enrichment of *Fusarium* antagonism in Sheriff pseudomonads, we determined BGC families that were differentially abundant in Heerup and Sheriff. Overall, there were not a difference in average number of BGCs in isolates from the two cultivars (two-sided Wilcoxon Rank Sum test, *p* > 0.05) (Fig 4A), but 28 of the 137 BGC families identified in the strain library were differentially abundant between the two cultivars (two-sided Wilcoxon Rank Sum test, fdr-adjusted *p* < 0.05) (Fig 4B). Nineteen and nine BGC families were enriched in Heerup and Sheriff, respectively, spanning 11 different product classes (Fig 4B). The only known BGC family differentially enriched between the two cultivars was BGC family 3049, an NRPS encoding a Pf-5 pyoverdine enriched in Sheriff (Table 2). Two CLPs were identified in the strain library, viscosin and lokisin, but they were not found to be differentially abundant between Heerup and Sheriff (Table 2).

**Figure 4.**
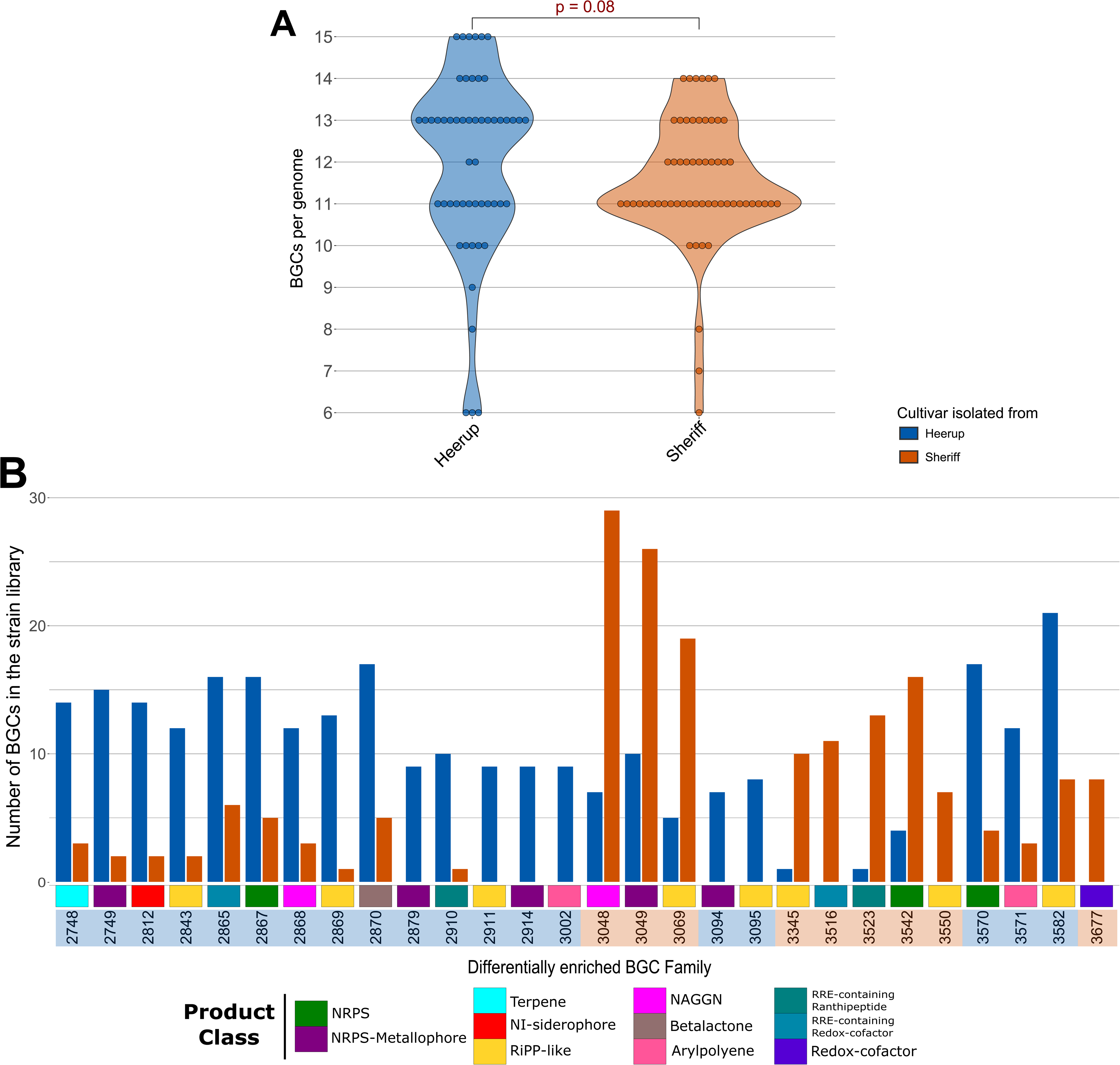
Biosynthetic potential in the Heerup and Sheriff strain libraries. A) Number of BGCs per genome of isolates cultured from Heerup and Sheriff. B) Differentially abundant BGC families found in Heerup- or Sheriff-associated pseudomonads. BGC families are color-coded along the x-axis by their predicted product class. The x-axis highlight box indicates if the BGC family is enriched in Heerup (blue) or Sheriff (orange).

Our next objective was to uncover the factors underpinning the differing abundance of BGC families within the Heerup and Sheriff *Pseudomonas* communities. First, we determined the effect of cultivar of origin on the number of BGCs per genome in each species (Figure 5A). Five species were found in both cultivars (*P. asgharzadehiana*, *P. simiae*, *P. neuropathica, P. siliginis*, and *P. brassicacearum*); however, only three of these species were found more than once on both cultivars (*P. asgharzadehiana*, *P. neuropathica*, and *P. brassicacearum*). Thus, statistical analysis was only done on these three species. The number of BGCs per genome was only affected by cultivar in the species *P. brassicacearum* (two-sided Wilcoxon Rank Sum test, *p* = 0.0140) (Figure 5A). Due to this unconvincing result of the number of BGCs per genome being dependent on cultivar of origin in only one species out of 22, we next analyzed the distribution of each differentially abundant BGC family across the strain collection (Figure 5B). This revealed that the variation in BGC family abundance was primarily driven by the enrichment or uniqueness of specific *Pseudomonas* species that were harboring many different BGC families to a single wheat cultivar.

**Figure 5.**
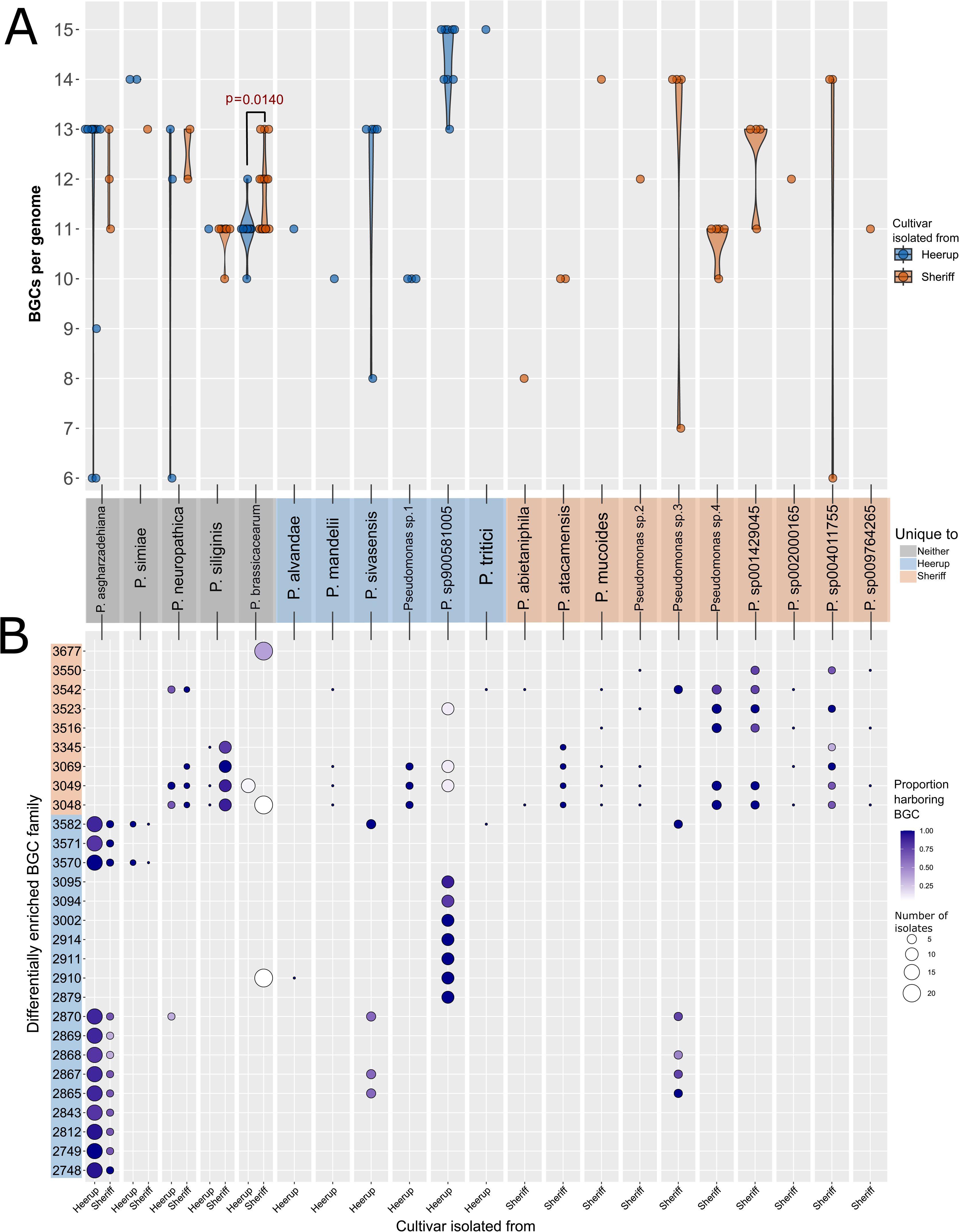
Distribution of BGCs in the species found in the Pseudomonas strain library. A) Number of BGCs per genome. True data points are overlaid the violin plot as filled circles. Significant differences in average BGCs per genome of a species between cultivars was calculated using the two-sided Wilcoxon Rank Sum test for the species P. brassicacearum, P. siliginis, and P. asgharzadehiana, and only p < 0.05 are displayed. B) Proportion of isolates of each species harboring the BGC family that is differentially enriched between Heerup and Sheriff. The x-axis highlight box represents if the Pseudomonas species is unique to Heerup (blue), Sheriff (orange), or found on both cultivars (grey). The y-axis highlight box represents if the BGC family is enriched in Heerup (blue) or Sheriff (orange).

### 3.5 Root scanning relates Sheriff *Pseudomonas* community to thinner root diameter

Culture-dependent and culture-independent 16S rRNA gene sequencing indicated that the Sheriff cultivar harbors a more diverse *Pseudomonas* population. To test the hypothesis that a more diverse *Pseudomonas* community would coincide with a thinner root diameter, we scanned the roots of four-week old Heerup and Sheriff plants and measured the root diameter of the system. Sheriff had a higher percentage of roots under 0.4 mm in diameter, while Heerup had a higher percentage of roots above 0.5 mm (Figure 6). This contributed to the thinner average root diameter of Sheriff (0.35 ± 0.03 mm) compared to Heerup (0.40 ± 0.03 mm) (two-sided Student T test, *p =* 0.0026).

**Figure 6.**
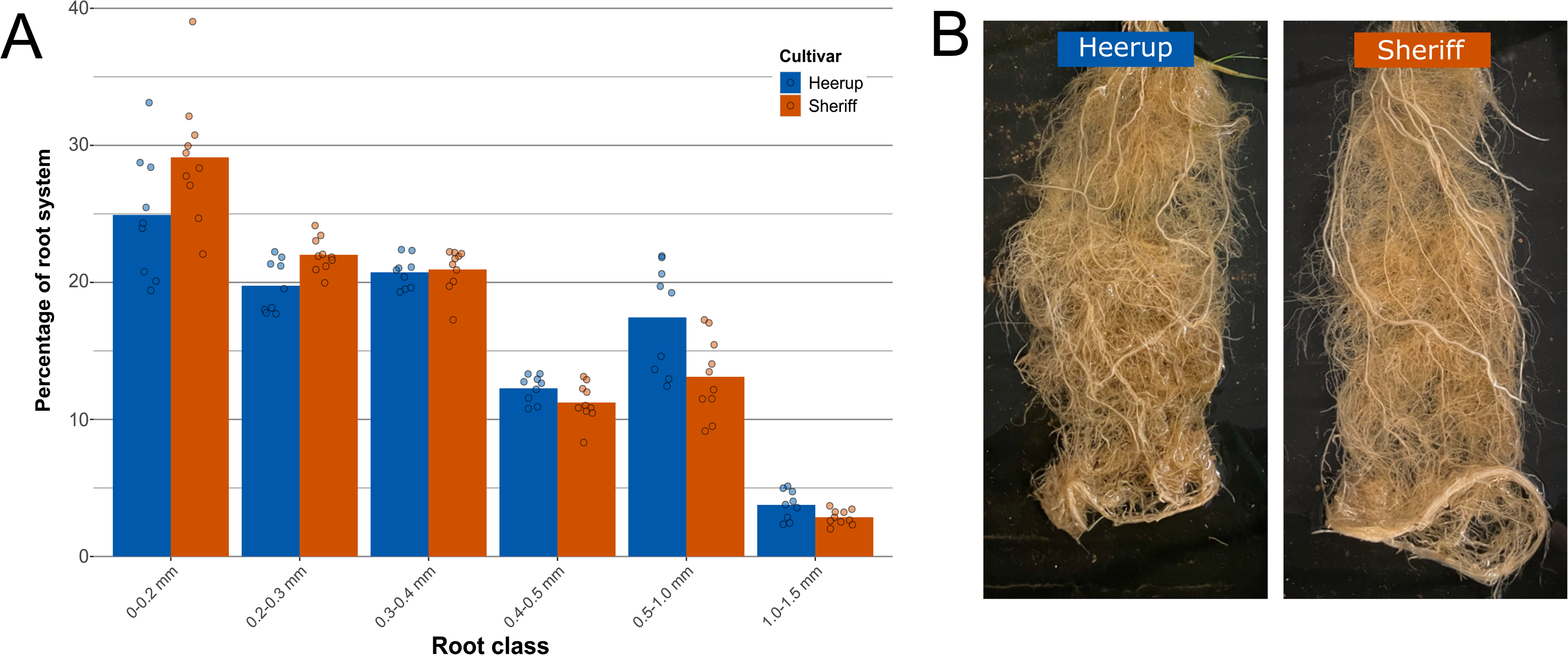
Root scanning on four-week old plants to measure root morphology parameters. A) Root system composition of Heerup and Sheriff divided into six root diameter ranges. The bar plot displays the mean percentage of root diameter in the root system of each cultivar. True data points are overlaid the bar plot as filled circles. B) Representative images of washed Heerup and Sheriff root systems after four weeks of growth.

## 4. Discussion

Our study provides an in-depth insight into the microdiversity of *Pseudomonas* communities in two commercially available modern cultivars of winter wheat with contrasting susceptibility to the fungal pathogen *F. culmorum.* With 373 isolates and 112 genomes, this is to our knowledge the largest study to characterize *Pseudomonas* microdiversity. Additionally, we provide a first glance into the *Pseudomonas* pangenome of two closely related modern cultivars, demonstrating that microbial microdiversity remains despite increasing homogeneity of agricultural systems (Elouafi, 2024).

In field-grown wheat rhizospheres, *Pseudomonas* are present in low abundances (Mauchline et al., 2015; Simonin et al., 2020), but their ability to dictate microbiome assembly (Garrido-Sanz et al., 2023; Getzke et al., 2023) and function (Hong et al., 2023; Lv et al., 2023) suggests their importance regardless. The results of our work indicate that the microdiversity of this genus leaves even more to be discovered. The taxonomic breadth in a single niche can be extensive, with our study revealing 22 species and 48 strains from 373 isolates as well as 70 ASVs from long-read 16S rRNA amplicon sequencing. Previous work in cocoyam and potato both identified seven phylogenetic groups from 134 and 69 *Pseudomonas* isolates, respectively (Oni et al., 2019; Pacheco-Moreno et al., 2021).

Interestingly, our low-abundance ASVs were different from our low-abundance isolates, indicating unique biases in both culturing and sequencing. Thus, we suggest that both approaches are needed to capture the full picture of *Pseudomonas* microdiversity in the rhizoplane.

In our first hypothesis, we predicted that the *F. culmorum* resistant cultivar, Sheriff, would have increased taxonomic diversity compared to the susceptible Heerup cultivar due to the link between microbial diversity and disease resistance (Hu et al., 2016; Li et al., 2023). Both culture-based and sequencing approaches indicated the preference of *Pseudomonas* species and strains to one cultivar or the other. These results agree with previous work that reports the ability of the plant to differentiate between phylogenetically similar bacteria (Thoms et al., 2023; Zhang and Kong, 2022). Culturable and *in silico* community analysis established that the Sheriff cultivar harbors a more taxonomically diverse *Pseudomonas* community, in support of our hypothesis. While not fully explored in this work, it may not be coincidental that the *Fusarium-*resistant cultivar Sheriff harbors two times the number of unique *Pseudomonas* strains on its roots compared to its susceptible relative, given the link between microbial diversity and plant health (Berg et al., 2017). Testing the protective ability of synthetic microbial communities of increasing intragenus complexity could support the idea of the beneficial effect of taxonomic microdiversity on plant health (Hu et al., 2016; Li et al., 2023).

Additionally in our first hypothesis, we predicted that increased taxonomic microdiversity in Sheriff pseudomonads would coincide with increased functional diversity, as *Pseudomonas* taxonomy is often related to functional abilities of the microbe (Lyng et al., 2024; Oni et al., 2019). While many COG20 functions were differentially abundant between the Heerup and Sheriff pseudomonads, they were equally distributed between the cultivars, giving no indication of increased functional diversity in one cultivar or the other. However, secondary metabolites biosynthesis, transport and catabolism was a COG20 category affected by cultivar. Thus, we analyzed the secondary metabolite biosynthetic potential of our strain collection in more detail. BGC families are one way to group architecturally similar BGCs that can be linked to a natural product chemotype (Navarro-Muñoz et al., 2020), and thus, may be used as a proxy for different potential functions based on secondary metabolites.

Interestingly, the less taxonomically diverse Heerup pseudomonads demonstrated increased biosynthetic functional diversity, boasting 19 enriched BGC families of 11 different product types while Sheriff only had nine families of five product types. The preferential colonization of the highly biosynthetic species *P. asgharzadehiana* on Heerup and the lack of biosynthetic genes in the many Sheriff-enriched species underpins this result. This goes against our preconceived hypothesis that increased taxonomic microdiversity results in increased functional microdiversity. Instead, few taxa were responsible for boosting biosynthetic diversity within the Heerup cultivar. This may be a conserved characteristic in soil, as bacterial taxonomic and functional diversity have previously been shown to be negatively correlated in this niche (Wang et al., 2023). Additional work in this area would elucidate the causes and consequences of such an inverse relationship and what this means for microdiversity as a beneficial trait in plant microbiomes.

Functional diversity within a microbial genus is only rarely examined between crop genotypes, and not yet between closely related modern cultivars (Oni et al., 2019; Pacheco-Moreno et al., 2024). We demonstrated the prevalence of microbial microdiversity in the absence of crop diversity by determining cultivar-dependent functional traits within the culturable *Pseudomonas* community. We found several instances of genomic loci enriched in one cultivar or the other. The distribution of BGC families also revealed cultivar-specific biosynthetic potential within the pseudomonads. In one case, it stemmed from cultivar-dependent interspecies microdiversity. The species *P. brassicacearum* carried BGC family 3677 only when it was isolated from the Sheriff cultivar. Interestingly, *P. brassicacearum* was also the only species where the average number of BGCs per genome was affected by cultivar, with Sheriff *P. brassicacearum* strains harboring more BGCs per genome in comparison to their Heerup counterparts. This may reflect an exciting ability of the plant to rely on bacterial secondary metabolites to structure its microbiome at the fine taxonomic scale. Previously, secondary metabolites have been implicated in driving plant distinction between closely-related microbes (Thoms et al., 2023) and used by *Streptomycetes* to carve a niche on the plant root (Nicolle et al., 2024). Deeper genome sequencing rhizosphere pseudomonads from additional wheat cultivars would confirm this interesting trend of cultivar-dependent interspecies microdiversity. It is also still up for debate why pseudomonads carrying these BGC families are found on their respective cultivar, especially since most of the BGCs in the strain library are unclassified beyond their product class. Further work in this area could resolve how crops recruit *Pseudomonas* producing certain metabolites to their microbiome, a mechanism previously suggested but poorly understood (Liu et al., 2021; Feng et al., 2023; Yang et al., 2023).

Our second hypothesis predicted that pseudomonads from the *F. culmorum*-resistant Sheriff cultivar would inhibit *F. culmorum* growth and harbor a greater abundance of genes capable of antagonizing *F. culmorum*. *Pseudomonas* are often implicated in disease suppression, even when grouped together at the genus level (Hong et al., 2023; Lv et al., 2023; Qiu et al., 2022). Thus, we examined antifungal activity as a microdiverse trait within the *Pseudomonas* genus. In support of our hypothesis, we found more *in vitro* antifungal activity against *F. culmorum* in the Sheriff isolates. Isolates that were positive for fungal inhibition also mapped to ASVs that were enriched in the Sheriff microbiome. In the pangenome, the BGC for the antagonistic metabolite 2,4-DAPG was found equally distributed between the two cultivars. Neither of the two CLPs found in the strain library, viscosin and lokisin, were enriched in either Heerup or Sheriff cultivar. In line with our hypothesis, we found a chitinase-encoding gene of the GH19 family enriched in Sheriff isolates. The GH19 family, originating from plants but also found in some bacteria including *Pseudomonas* (Ren et al., 2022), are notable for their activity against *Fusarium* (García-Fraga et al., 2015). We also found a predicted *P. fluorescens*-type pyoverdine BGC that was enriched in Sheriff pseudomonads. Bacterial siderophores empower phytopathogen control via competition for iron (Gu et al., 2020) and have been long documented as a *Pseudomonas* bioactive molecule against *Fusarium* (Kloepper et al., 1980). In agreement with our observations, siderophore-mediated iron uptake was previously shown to be a lineage-specific trait in *Pseudomonas* (Shalev et al., 2021). Thus, the specific *Pseudomonas* traits present in a microbiome may be more important than the abundance of the *Pseudomonas* genus as a whole in the context of plant health. Future efforts to analyze the plant holobiont for improved health should therefore be focused on function, rather than taxonomy.

Our last hypothesis predicted that the wheat cultivar with the higher taxonomically diverse microbiome would have a thinner average root diameter, and in support of this hypothesis, we saw that the more diverse Sheriff cultivar had a thinner average root diameter and a higher proportion of thinner roots compared to Heerup. This contributes to the growing body of literature that describes a more diverse microbiome in both woody and herbaceous plant species which have thinner roots or a root system dominated by fine roots (Saleem et al., 2018; Pervaiz et al., 2020; Luo et al., 2021; Zai et al., 2021; Fleishman et al., 2023). Nevertheless, it is not possible at this time to attribute the differences in *Pseudomonas* communities to the root morphology of the cultivars, since the cultivars are genetically different. Other factors including root exudation and other plant genes are certainly at play (Sasse et al., 2018; Song et al., 2021). Still, root diameter and root morphology as a whole might play a previously unexplored role in microbial community composition, as we have suggested previously (Herms et al., 2022), and deserve further evaluation.

## 5. Conclusion

The rhizosphere microbiome is a key determinant of plant health, and ongoing research aims to unravel the secrets to utilizing beneficial plant-microbe interactions to promote sustainable agriculture. As agricultural systems become increasingly homogenous, it is important to understand how closely related modern cultivars can still differ in their microbiomes. In this work, we demonstrate that the status quo of grouping together highly diverse genera, such as *Pseudomonas*, in 16s rRNA analyses overlooks crucial differences in microbial functional activity (Chiniquy et al., 2021; Jaspers and Overmann, 2004). Failing to consider the importance of microdiversity in shaping the composition and function of the rhizosphere microbiome conceals the true potential of root-associated microbial communities and should be a focus area in future plant holobiont research. We suggest that microdiversity should be explored as a potential metric for the health and productivity of plant microbiomes, but the relationship between taxonomy and function remains a confounding factor on the mechanism underpinning taxonomic diversity and pathogen suppression. In addition to providing novel insight into the microbial microdiversity of elite crop cultivars, our unique collection of cultivar-specific *Pseudomonas* isolates can act as a tool to further dissect high-resolution plant-microbe interactions. Exploration of these cultivar-dependent genomic loci may reveal insights regarding the specificity of plant-microbe interactions at the species level. Ultimately, unraveling this interaction can allow for designer crops that use microbial metabolites for increased resilience in the face of challenging growing conditions.

## CREDIT authorship statement

Courtney Horn Herms: Conceptualization, Data Curation, Formal analysis, Investigation, Methodology, Project administration, Software, Visualization, Writing - Original Draft, Writing - Review & Editing. Rosanna Catherine Hennessy: Conceptualization, Investigation, Methodology, Supervision, Writing - Review & Editing. Frederik Bak: Conceptualization, Data curation, Formal analysis, Methodology, Software, Visualization, Writing - Review & Editing. Ying Guan: Investigation, Methodology, Writing - Review & Editing. Patrick Denis Browne: Data curation, Formal analysis, Methodology, Software, Visualization, Writing - Original Draft, Writing - Review & Editing. Tue Kjærgaard Nielsen: Data curation, Formal analysis, Methodology, Software, Writing - Review & Editing. Lars Hestbjerg Hansen: Methodology, Resources, Writing - Review & Editing. Dorte Bodin Dresbøll: Methodology, Resources, Writing - Review & Editing. Mette Haubjerg Nicolaisen: Conceptualization, Funding acquisition, Methodology, Project administration, Resources, Supervision, Writing - Review & Editing.

## Conflict of Interests

The authors report no conflicts of interest.

## Acknowledgements

We thank Dorette Müller-Stöver and Marie Louise Bornø, University of Copenhagen, for supplying experimental soils. The *Fusarium culmorum* strain was kindly provided by Birgit Jensen, University of Copenhagen. Thanks to our group members Kitzia Yashvelt Molina Zamudio and Dorthe Thybo Ganzhorn for their support with the sampling.

## Funding

This work was supported by the Novo Nordisk Foundation [grant number NNF19SA0059360].

## Abbreviations

ASV: amplicon sequence variant
BGCs: Biosynthetic gene clusters
CLPs: cyclic lipopeptides
CSV: culture sequence variant
NRPS: non-ribosomal peptide synthetase
RiPP: ribosomally synthesized and post-translationally modified peptide

---

**Methods Figure 1.**
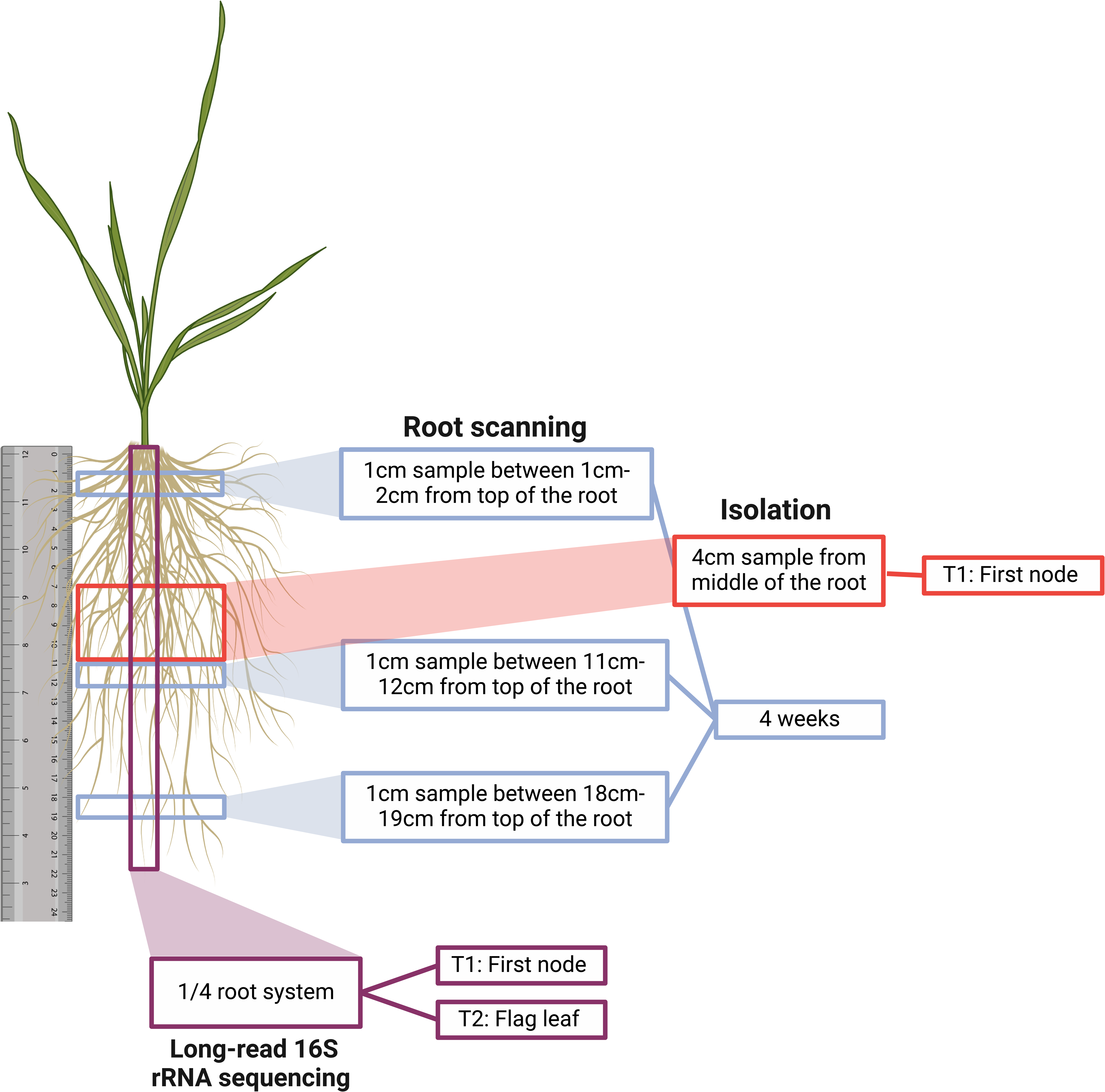
Sampling for Pseudomonas isolation, long-read 16S rRNA amplicon sequencing, and root scanning occurred along different root axes as well as across different time points. Created with Biorender.com.

## Notes

https://www.ncbi.nlm.nih.gov/sra/PRJNA1108081

### Competing Interest Statement

The authors have declared no competing interest.

### Summary of Updates

Introduction and Discussion re-written to improve understanding and clarity. Minor figure updates. No changes to data analysis or interpretation.

