## Supplementary figures and images for "*Pseudomonas* taxonomic and functional microdiversity in the wheat rhizosphere is cultivar-dependent and links to disease resistance profile and root diameter"

### Supplemental Figure 1

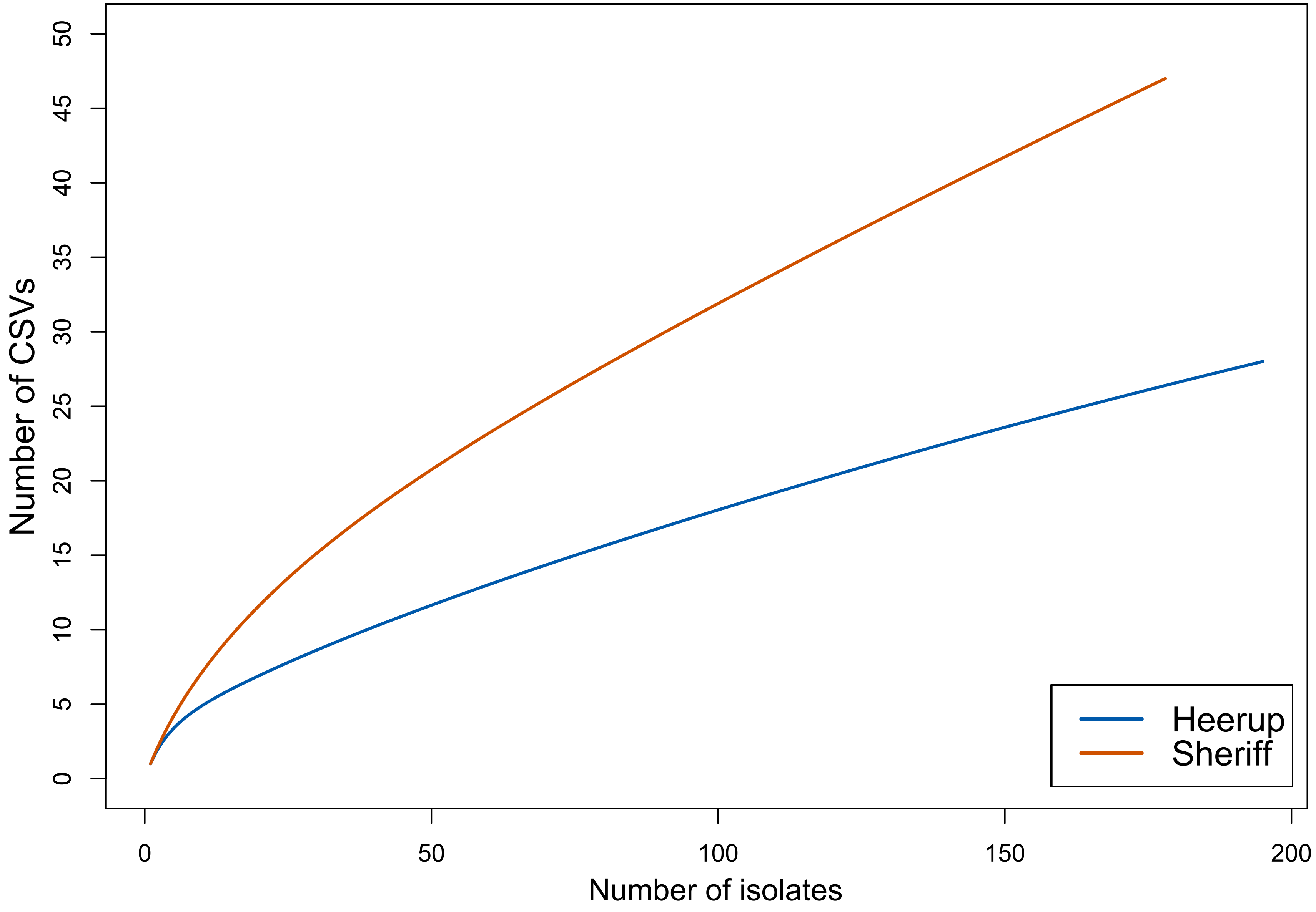

### Supplemental Figure 2

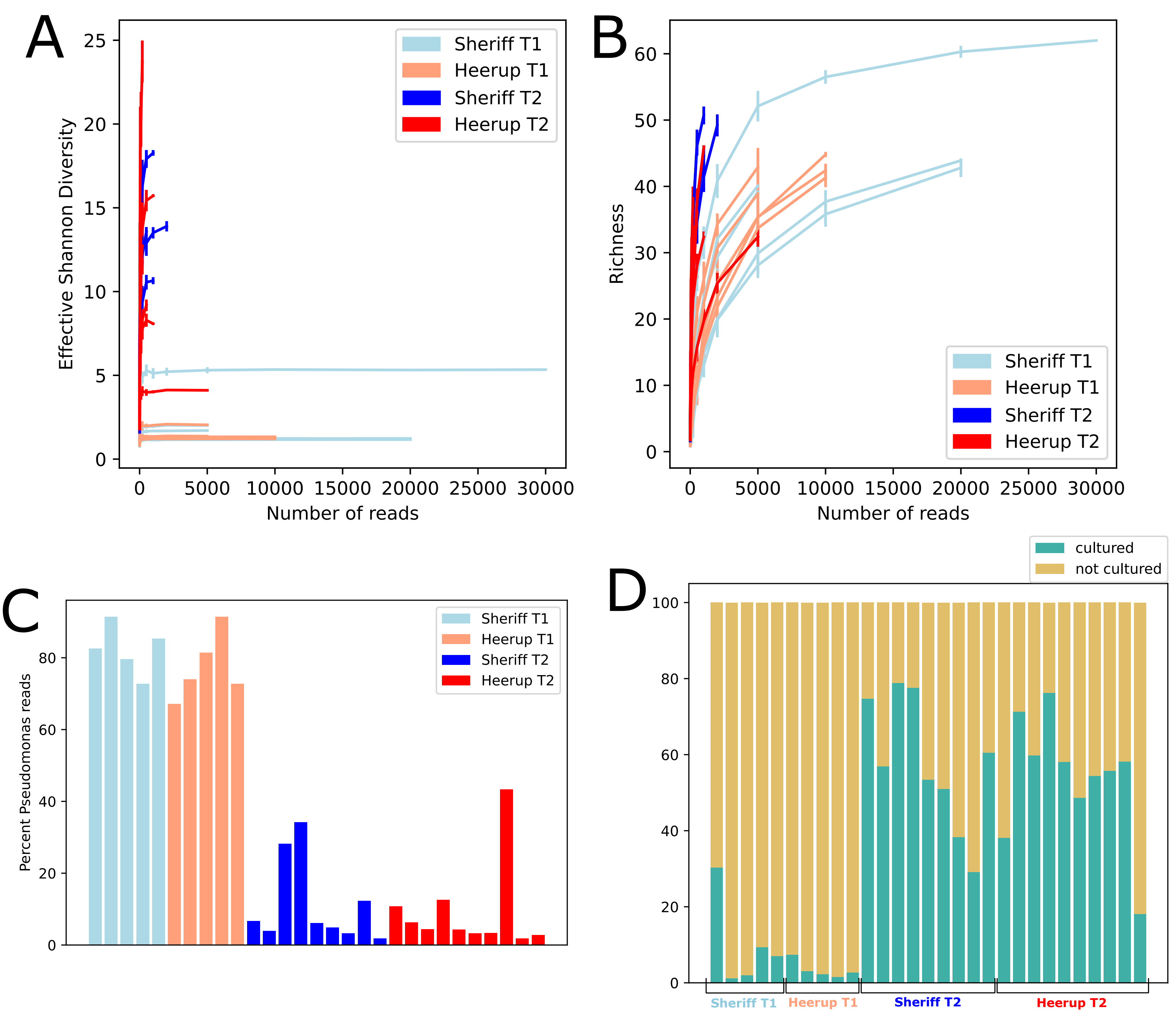
