## Supplemental File for "*Pseudomonas* taxonomic and functional microdiversity in the wheat rhizosphere is cultivar-dependent and links to disease resistance profile and root diameter"

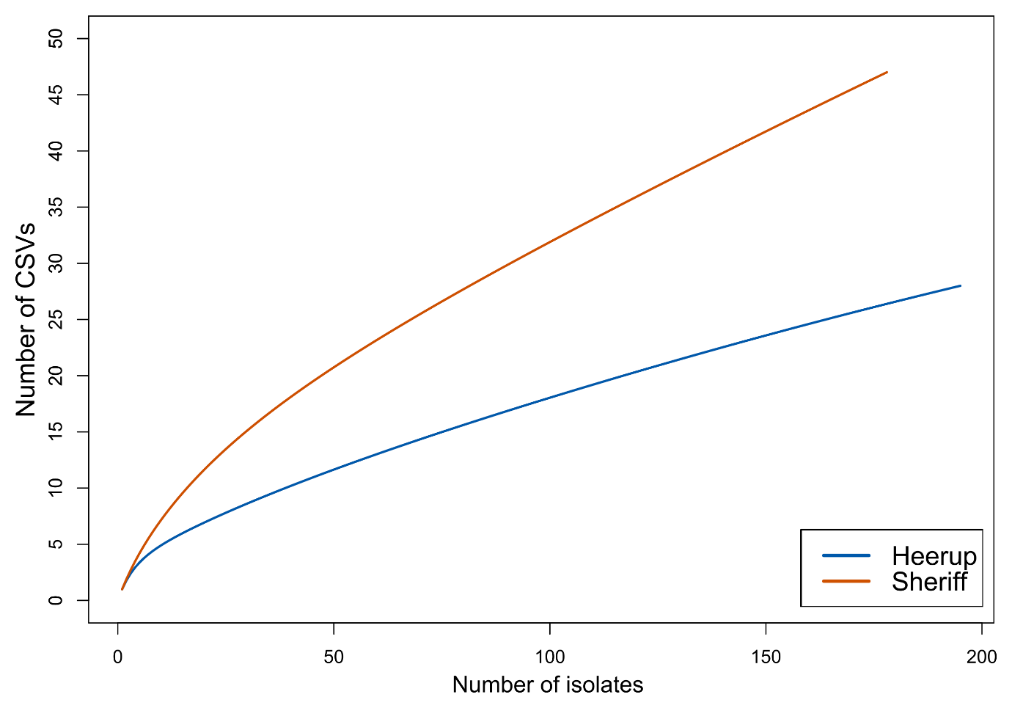

**Figure S1. Rarefaction curve based on CSV identification during sampling of *Pseudomonas* isolates from the two cultivars of wheat.**

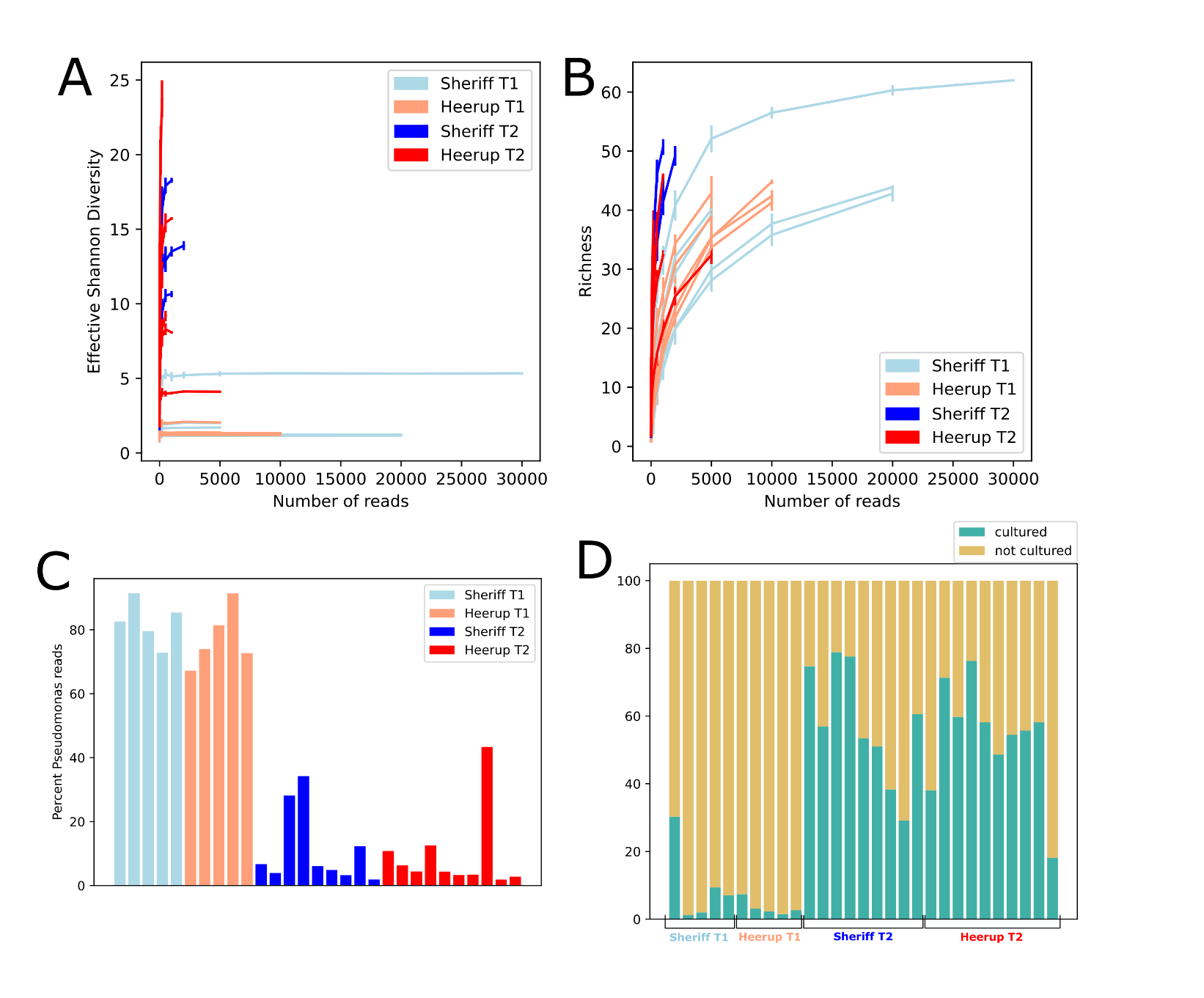

**Figure S2. Sequencing results from long-read 16S rRNA amplicon sequencing using Nanopore technology across 880-970 bp of the 16S rRNA gene.** A) Rarefaction plot for effective Shannon diversity and B) richness. C) Percent of reads per sample that were identified as *Pseudomonas*. D) Percent of *Pseudomonas* reads per sample that mapped to cultured isolates or uncultured sequences.

**Table S1. Counts of *Pseudomonas* found during each method used in the study, based on the various taxonomic levels and their definition based on percent identity across the sequence.**

**Table S2: COG categories and functions for gene clusters identified by Anvi'o as enriched in a *Pseudomonas* species or strain isolated uniquely from one wheat cultivar.**

**Table S3: COG20 categories in the *Pseudomonas* pangenome identified by Anvi'o as enriched in one wheat cultivar.**

**Table S4: COG20 functions in the *Pseudomonas* pangenome identified by Anvi'o as enriched in one wheat cultivar.**
